# Tetraspanin Cd9b plays a role in fertility in zebrafish

**DOI:** 10.1101/2021.11.25.469903

**Authors:** S Greaves, KS Marsay, PN Monk, H Roehl, LJ Partridge

## Abstract

In mice, CD9 expression on the egg is required for efficient sperm-egg fusion and no effects on ovulation or male fertility are observed in CD9 null animals. Here we show that *cd9b* KO zebrafish also appear to have fertility defects. In contrast to mice, fewer eggs were laid by *cd9b* KO zebrafish pairs and, of the eggs laid, a lower percentage were fertilised. *In vitro* fertilization (IVF) was used to demonstrate that factors such as courting behaviour, adult size and mate choice were not causing the unexpected decrease in clutch size. The decrease in egg numbers could be rescued by exchanging either *cd9b* KO partner, male or female, for a wildtype (WT) partner. However, the fertilisation defect could only be rescued by crossing a *cd9b* KO female with a WT male. Our results indicate that Cd9b has several roles in fish fertility, affecting both clutch size and egg fertilisation.

**Highlights:**

1. *cd9b* mutant pairs lay fewer eggs and of the eggs laid, fewer are fertilised.
2. Mutation in *cd9b* does not affect primordial germ cell number or migration.
3. Reduction in number of eggs and fertility is not due to courting behaviour, fish size or mate choice.
4. *cd9b* mutation in either gender affects the number of eggs laid.
5. Mutation of *cd9b* in the males affects fertilisation efficiency but mutation of *cd9b* in the females does not.

## Introduction

CD9 is a member of the tetraspanin superfamily of proteins that function as organisers of other membrane proteins (van Deventer et al., 2017). CD9 is involved in a wide range of cell functions, including adhesion, motility, signalling and cell fusion (Reyes et al., 2018). Knockout (KO) of CD9 in female mice results in infertility due to a defect in sperm/oocyte fusion (reviewed in (Jankovicova et al., 2015)). In a mechanism that appears conserved in mammals, CD9 is suggested to partner Juno, the egg receptor for sperm ligand Izumo1, thus facilitating the formation of adhesion sites prior to fusion (Chalbi et al., 2014). CD9 concentrates at the interaction site on the oocyte in response to sperm oscillations immediately before fusion (Ravaux et al., 2018). Whilst male CD9 KO mice appear fertile, CD9 is expressed on mouse sperm and male germline stem cells (Ito et al., 2010, Frolikova et al., 2018) and is present at various stages of spermatogenesis, suggesting a role in this process (Zohni et al., 2012).

Tetraspanins are widely expressed in teleosts (Cao and Tan, 2018) but there are no reports of roles in fish fertility. Several tetraspanins have roles during zebrafish development, including pigment cell interactions (Inoue et al., 2014), hatching (Trikic et al., 2011), vascularisation (Li et al., 2012; Li et al 2020), migrasome formation (Jiang et al., 2019) and primordium migration (Marsay et al., 2021). However, the role of Cd9 in fertilisation has not yet been investigated. There are two paralogues of *cd9* in zebrafish (*cd9a* and *cd9b*), which have 63% amino acid identity and similar mRNA expression patterns (Marsay et al., 2021).

In this report, we investigate the role of Cd9 in zebrafish fertility. Two zebrafish *cd9b* alleles were used, and homozygous incrosses of both alleles exhibited defects in fertilisation rates. The number of eggs produced per female (clutch size) was also significantly reduced. The defect in fertilisation was not further exacerbated by the additional KO of the paralogue, *cd9a*. Reduced clutch size could be rescued by crossing either *cd9b* KO male or female fish with a WT partner. In contrast, reduced fertilisation could only be rescued by crossing a KO female with a WT male. Our results indicate that Cd9 plays a more complex role in fish fertility than in mammals, with effects on both male and female fertility.

## Methods and materials

### Zebrafish Maintenance

Adult wildtype zebrafish (WT) and *cd9a/b/dKO* mutants were housed and bred in a regulated 14:10 hour light: dark cycle under UK Home Office project licence 40/3459 in Bateson Centre aquaria at the University of Sheffield or project licence IACUC 140924 in the Singapore IMCB zebrafish facility. Zebrafish were raised under the standard conditions at 28°C (Nüsslein-Volhard and Dahm, 2002).

### Zebrafish mutant production

*cd9b* mutants were created from WT embryos using transcription activator-like effector nucleases (TALEN) and maintained on an WT background. TALENs (ZGene Biotech Inc., Taiwan) were provided in a pZGB4L vector, targeting the *cd9b* sequence 5’ ttgctctttatcttca 3’. Two frameshift mutants, c.46del (*cd9b^is16^* allele) or c.42_49del (*cd9b^pg15^* allele) were selected that caused premature termination just after the first transmembrane domain (Fig S1) (previously described by Marsay et al., 2021). *cd9a* mutants were created by Marsay et al., 2021 using CRISPR/Cas9. An indel mutation deleting 4bp and inserting 8bp (c.180_187delinsTCGCTATTGTAT; *cd9a^la61^*) generated a frameshift mutation resulting in a premature stop codon in exon 3, which was predicted to truncate the protein before the large extracellular domain. *cd9* dKO mutants were created by injecting the *cd9a* gRNA and Cas9 RNA into *cd9b^pg15^* embryos. These fish were screened for germline transmission by sequencing and backcrossed to *cd9b^pg15^* mutants. Heterozygous fish of the same genotype were incrossed and adult F2 fish were genotyped to identify homozygous *cd9b^pg15^*; *cd9a^la61^* (*cd9* dKO).

### Embryo collection and analysis

Adult zebrafish male/female pairs were placed in plastic breeding tanks overnight, separated by a divider. The divider was removed the following morning after the lights came on and spontaneous spawning occurred. Embryos were collected every 20 min and collection time was recorded. Zebrafish pairs were allowed to spawn until no more embryos were produced and the number of embryos produced by each pair was recorded. Dead eggs (opaque eggs) were counted and removed on collection and fertilisation was assessed four hours post collection. Embryos that presented a well-developed blastodisc were counted as fertilised.

### Probe synthesis for *in situ* hybridisation

*vasa* cDNA was provided by H. Knaut (NYU Medical Center and School of Medicine, USA) in a pBS+ cloning vector. The vector conferred ampicillin resistance and contained M13 primer binding sites flanking the *vasa* cDNA. *vasa* cDNA containing plasmid was transformed into NEB 10-beta competent *E.coli*, and purified using a Miniprep kit (Qiagen, UK). The DNA template for the *vasa* RNA probe was then produced using a standard PCR protocol with M13 primers (Forward: 5’gtaaaacgcggccagt3’, Reverse: 5’ggaaacagctatgaccatg 3’), and purified using a 50 kDal centrifugal filter unit (Amicon, UK). Anti-sense RNA probes were transcribed from the DNA template using digoxigenin (DIG)-11-UTP Labelling Mix (Roche, UK), cleaned using spin filters (Sigma-Aldrich) and eluted into RNA-later (Sigma-Aldrich, UK) before storing at −20°C.

### *In situ* hybridisation

Embryos were raised at 28°C in petri dishes containing E3 solution. The E3 was changed daily and any dead embryos removed. At 30-32 hours post fertilisation (hpf), embryos were anaesthetised using tricaine, dechorionated and then fixed using 4 % (w/v) paraformaldehyde (PFA; Sigma-Aldrich, UK) in PBS. The fixed embryos were left overnight at 4°C in 4 % PFA before being washed twice with PBS/0.05 % (v/v) Tween 20 (PBST) the following morning. Embryos were then put through a MeOH/PBS series using 30 %, 60 % and 100 % (v/v) MeOH before being stored in 100% MeOH (Sigma-Aldrich) at −20°C. *In situ* hybridisation was carried out as described (Thisse and Thisse, 2008), except for the embryo digestion with proteinase K, for which 30-32 hpf embryos were digested with 10 mg/ml proteinase K at 20°C for 22 min. The protocol was performed with embryos in 1.5 ml microfuge tubes for the first two days, after which they were placed in 12-well plates for staining before transferring back to microfuge tubes for storage. Stained embryos were stored in the dark in 80% (v/v) glycerol.

### Primordial germ cell (PGC) assays

PGC were stained using a *vasa in situ* hybridisation and then embryos were imaged in 80% glycerol using a microscope mounted camera and a 5× or 10× objective. The number of PGCs was counted across the whole embryo and PGC migration was analysed by measuring the distance between the most anterior and posterior PGCs, with the measurement following the body axis. Measurements were taken using Image J software.

### *In vitro* fertilisation (IVF)

Adult zebrafish were paired, as described above, 4 days before the IVF procedure and then transferred back to their normal tanks. The fish were then paired again in the afternoon before the IVF procedure. The following morning, fish of the same genotype were placed together in larger tanks as zebrafish will not normally lay when grouped. Individual fish were then anaesthetized using tricaine and dried before gamete extraction. Sperm was extracted from male fish using suction through 10 μl capillaries (Hirschmann Laborgeräte GmbH, Germany), whereas females were gently pressed on the abdomen to release eggs. Gametes from a single pair of individuals were combined and incubated for 30 sec before adding 750 μl aquarium water and incubating for a further 2 min. 9 ml of aquarium water was then added and the gametes incubated for 4 hr at 28°C. The numbers of fertilised and unfertilised eggs were then assessed. Dead eggs were immediately discarded after extraction from the females and therefore not included in the analysis.

### Statistics

Data distribution was first assessed for normality using a D’Agostino-Pearson omnibus K2 normality test on the experimental residuals, as well as creating a histogram of residuals. For normally distributed data, an ANOVA with Dunnet’s or Holms-Sidak multiple comparisons tests were used. For non-normally distributed data non-parametric tests, the Mann-Whitney U test or Kruskal-Wallis with Dunn’s multiple comparisons test, were used.

## Results and Discussion

To test the involvement of Cd9b in zebrafish fertility, we used two alleles of Cd9b, *cd9b^is16^* and *cd9b^pg15^*. The homozygous mutant KO fish appeared to develop normally. However, when incrossed, both the number of eggs per clutch (Fig 1A) and the fertilisation rate of the eggs produced were significantly reduced compared to WT (Fig 1B). Although fertility was reduced in both alleles, the extent of reduction differed dramatically between the two alleles (Fig 1B). The loss of fecundity in both alleles was surprising because the KO of CD9 in mice affects only the fertilisation of ova and not their production (Le Naour et al., 2000). When the fate of the zebrafish eggs was analysed in more detail, the *cd9b^is16^* KOs produced a significantly higher percentage of eggs that were dead at the time of embryo collection (Fig 1C). Dead eggs are opaque and are easily identified. However, this was not observed in the *cd9b^pg15^* KO line, which produced a significant number of live but unfertilised eggs. (Embryos that presented a well-developed blastodisc after 3 hours were counted as fertilised). This suggests that decreased survival of the eggs is not the only reason for decreased fertility. To determine if deletion of both paralogs would result in a complete loss of fertility as seen in the CD9 KO mouse, *cd9a* was knocked out in the *cd9b^pg15^* background. Interestingly, the fertilisation rate of the double KO line was very similar to the *cd9b^pg15^* KO line (Fig 1D), suggesting that only Cd9b is involved in egg production and fertilisation.

**Figure 1.**
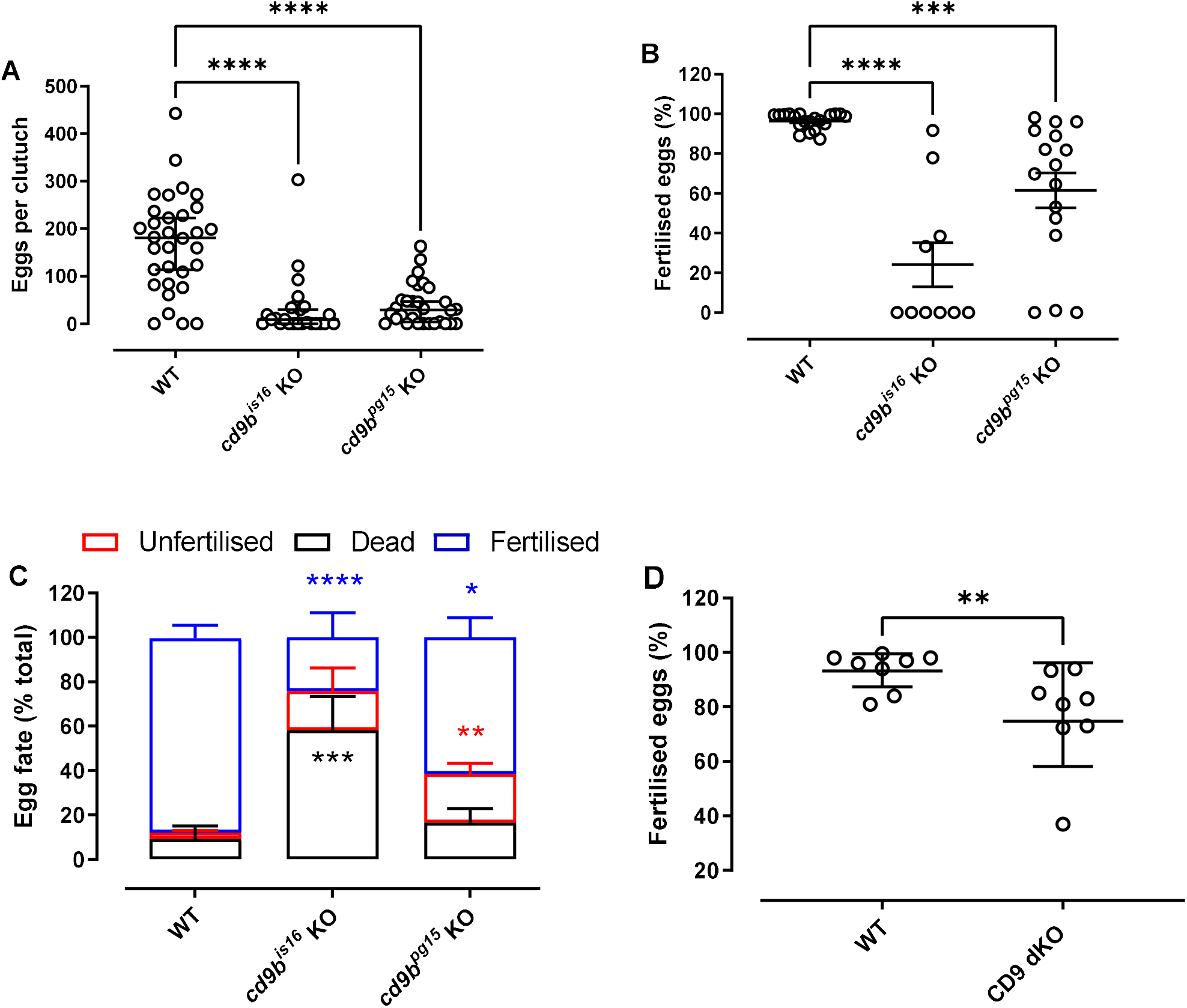
*cd9b* mutant pairs lay smaller clutches of eggs with significantly fewer fertilised eggs. A significant decrease in the total number of eggs laid per pair, including fertilised, dead and unfertilised eggs, is seen with *cd9b* mutant pairs (A). The percentage of fertilised eggs is significantly decreased in clutches from *cd9b* mutant pairs (B). Given the significant decrease in fertilised eggs seen in (B), it can be inferred that there is a significant increase in the percentage of dead and unfertilised eggs per clutch from *cd9b* mutants. This increase is apparent in (C), where the average breakdown of clutches from WT and *cd9b* mutant pairs is shown. The additional loss of *cd9a* in *cd9* dKOs did not result in a further decrease in the fertilisation rate (D). (A), (B) and (C) represent pooled data from 5 experimental repeats. A Kruskal-Wallis test with Dunn’s multiple comparisons test where p<0.05 was performed on (A). An ANOVA with Dunnett’s multiple comparisons test was performed on (B, C) after removal of definitive outliers (3 outliers, ROUT, q=0.1) to give normally distributed data (p<0.05). n=minimum 25 pairs (A), minimum 10 pairs (B, C) per genotype. Only pairs that laid eggs were counted in (B) and (C), whereas all pairs were counted in (A). (D) represents 8 separate matings, with a Mann-Whitney U test of significance. Significance of difference from WT control: **** p<0.001; *** p<0.005; ** p<0.01; * p<0.05; ns=not significant.

We then investigated primordial germ cell (PGC) behaviour, to determine if reduced numbers or a delayed migration could result in lowered egg production (Wong and Collodi, 2013). However, both *cd9b* KO lines have the same number of PGC as WT fish (Fig 2A) and migration to the gonadal ridge during development was not altered in *cd9b* mutants (Fig 2B).

**Figure 2.**
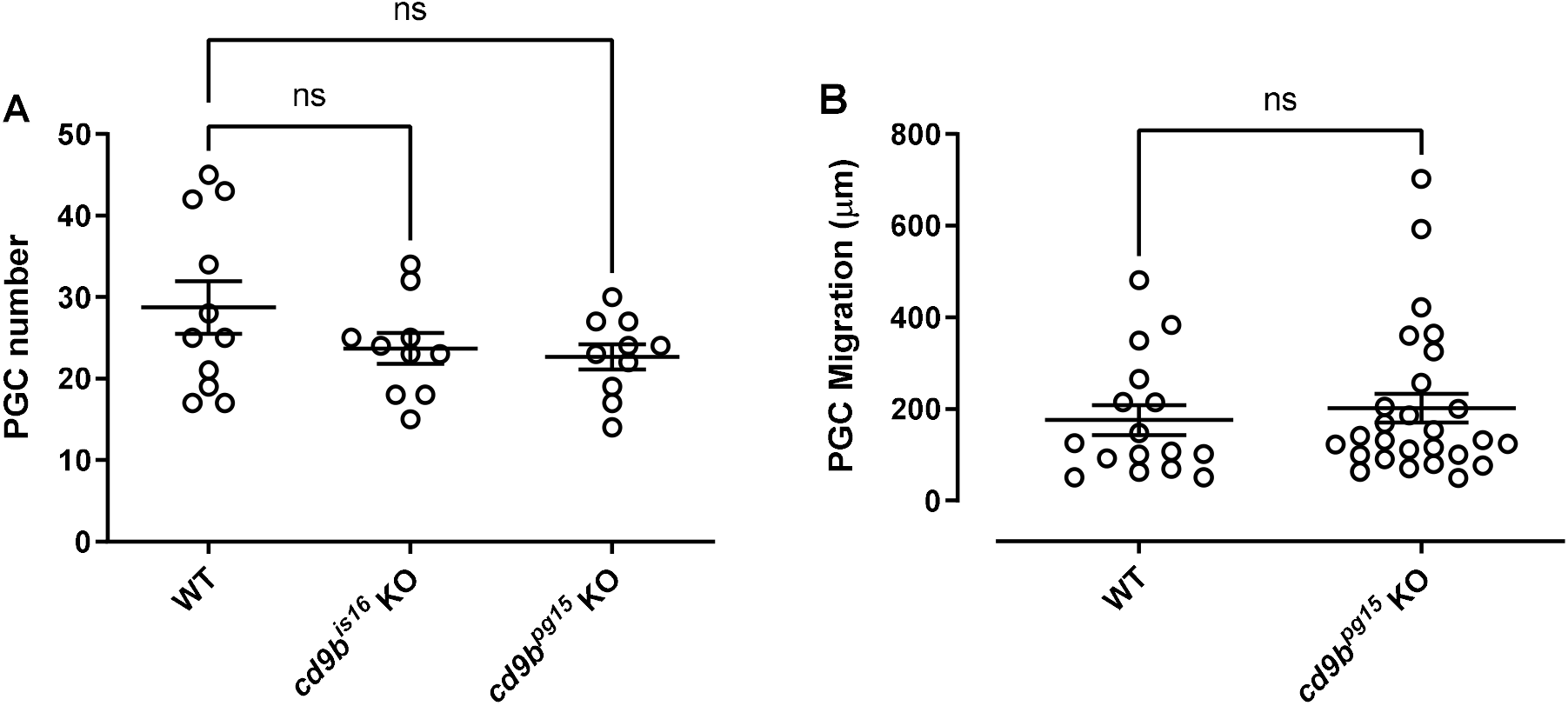
*cd9b* homozygous mutants have the same number of PGC per embryo at 32 hpf and migration to the gonadal ridge is not affected in *cd9b^pg15^* mutants at 30 hpf. (A) There is no significant difference in the number of PGCs in WT and *cd9b* mutant embryos at 32 hpf. PGCs were visualised using a *vasa* ISH on 32 hpf embryos and analysed using an ANOVA with Dunnett’s multiple comparisons test on n=minimum 10 embryos per genotype. (B) *vasa* ISH on 30 hpf WT and *cd9b^pg15^* embryos. PGC migration efficiency was analysed by looking at the distance between the most anterior and posterior PGC, which should have reached the gonadal ridge by 30 hpf. No significant difference was seen between WT and *cd9b* mutants as shown by a Mann-Whitney U test. n=at least 13 individual embryos.

It is possible that egg production and fertilisation in zebrafish are affected by mating behaviour, as observed previously (reviewed in (Nasiadka and Clark, 2012)). To investigate this, we attempted to fertilise eggs using IVF techniques. We found that numbers of fertilised embryos obtained from female *cd9b* mutants was significantly lower than WT (Fig 3A). Fertilisation rates using sperm from *cd9b* KO males were also significantly reduced (Fig 3B). This suggests the reductions in clutch size and fertilisation in *cd9b* KO mutants are not due to size differences between fish when pairing, or courtship behaviours such as chasing.

**Figure 3.**
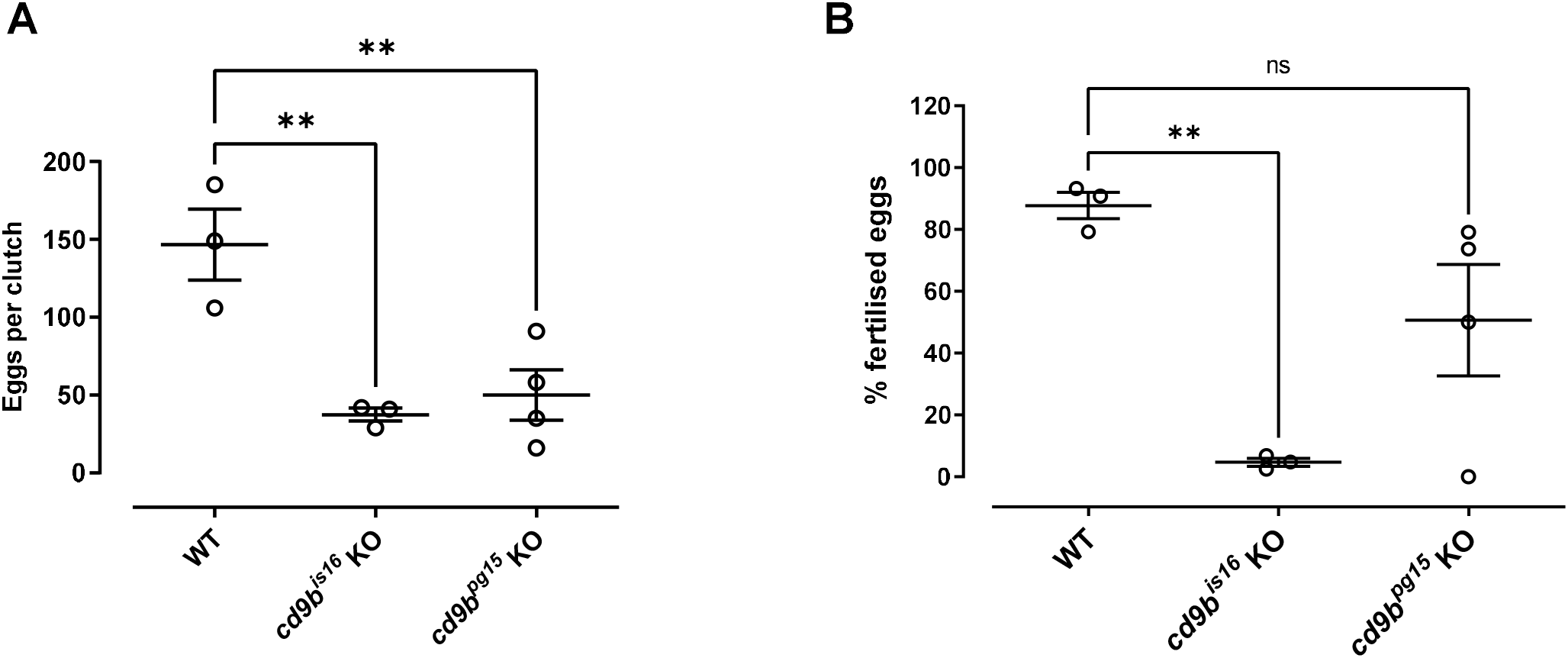
Clutches from *cd9b* mutant pairs, produced by IVF, have a decreased clutch size and lower percentage of eggs fertilised per clutch. The number of eggs is significantly decreased in clutches from *cd9b* mutant pairs (A). Of the eggs laid, a significantly lower percentage were fertilised by *cd9b* mutant males (B). Dead eggs were immediately discarded during the IVF protocol and therefore (A) and (B) represent only fertilised and unfertilised eggs. n=minimum 3 pairs per genotype. ANOVA with Holm-Sidak multiple comparisons test was carried out on the original data from (A) and on arcsine transformed data in (B). The Holm-Sidak post-hoc test was chosen as it has more power than Dunnett’s multiple comparisons test. Significance of difference from WT control: ** p<0.01; * p<0.05; ns=not significant.

These data suggest that the defects caused by *cd9b* KO may not just be due to effects on female fish but may also involve aspects of male fertility. To investigate this, we measured clutch size and fertilisation rates using a matrix of crossings. As found previously, mutant females crossed with mutant males of either KO line had decreased clutch size and fertilisation rates (Fig 4A, B). Crossing *cd9b* mutants of either gender with WT fish produced normal clutch sizes (Fig 4A), showing that this phenotype can be rescued by both male and female WT fish. In contrast, the defect in the percentage of eggs fertilised was only rescued when the *cd9b* mutant male was substituted for a WT male. This suggests that the reduction in fertilisation seen in *cd9b* mutant pairs is due solely to a difference in the mutant male.

**Figure 4.**
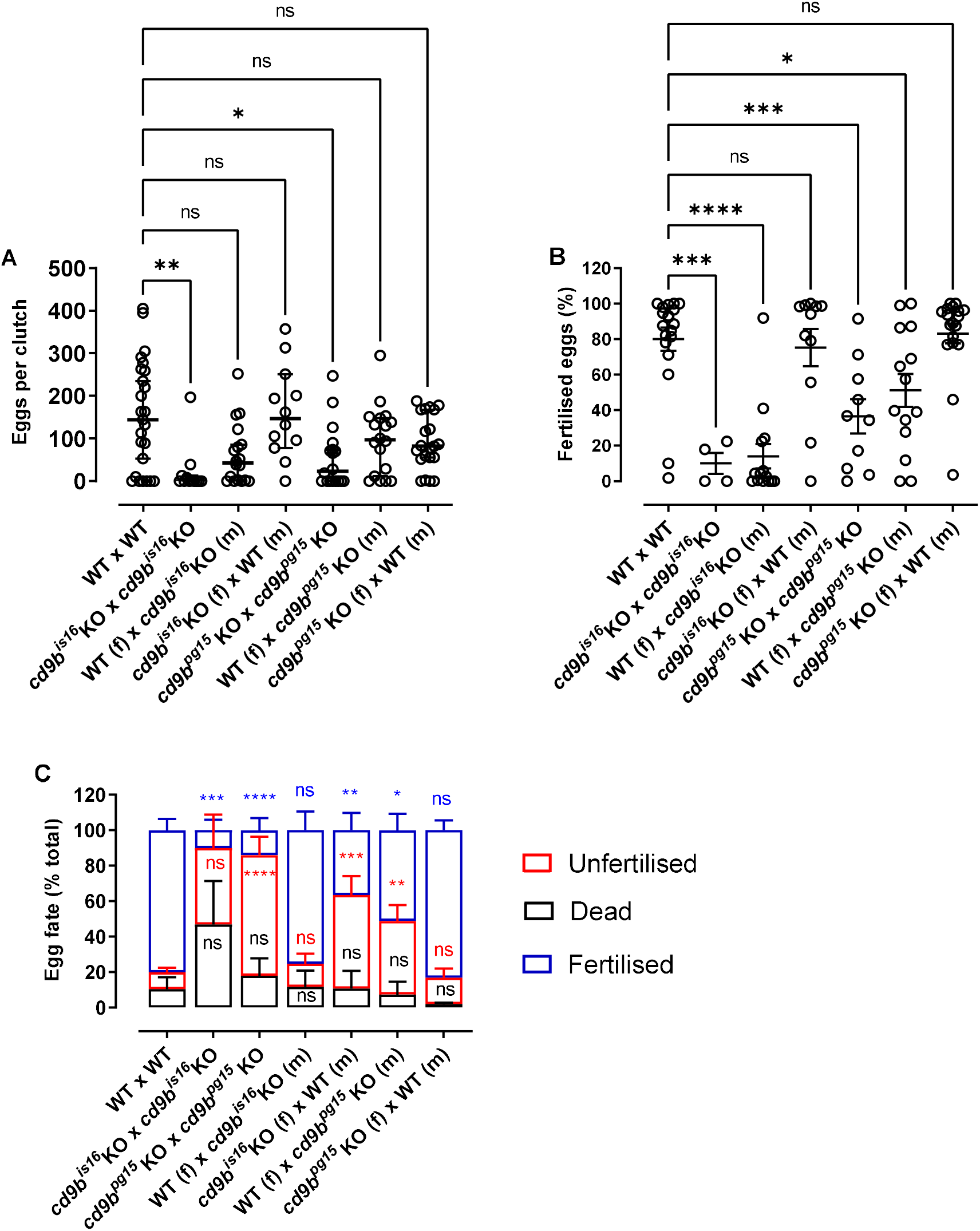
Clutch size can be rescued by substitution of either *cd9b* mutant partner for an WT, however, fertilisation is only rescued when the *cd9b* KO male is substituted for an WT male. Clutch size can be rescued by substitution of a *cd9b* mutant partner, male (m) or female (f), with an WT (A). A Kruskal-Wallis test with Dunn’s multiple comparisons test was carried out on the pooled data from three experimental repeats. n=minimum 12 pairs per cross. While clutch size can be partially or fully rescued by substitution of either *cd9b* mutant partner, the percentage of fertilised eggs appears to only return to WT levels upon substitution of a *cd9b* mutant male with an WT male (B). An ANOVA with Dunnett’s multiple comparisons test was carried out on arcsine transformed data pooled from three experiments. The average percentage of fertilised, dead and unfertilised eggs per clutch is shown in (C). n=minimum 10 pairs per cross for (B) and (C), except *cd9b^is16^* × *cd9b^is16^* where only 4 of 12 pairs laid. Significance of difference from WT control: **** p<0.001; *** p<0.005; ** p<0.01; * p<0.05; ns=not significant.

In conclusion, as with CD9 KO mice, *cd9b* homozygous mutant zebrafish showed fertility defects. It was found that *cd9b* KO zebrafish pairs laid decreased numbers of eggs and *cd9b* KO males had severely reduced fertility.

