## Supplementary figures for "Tetraspanin Cd9b plays a role in fertility in zebrafish"

### Supporting information

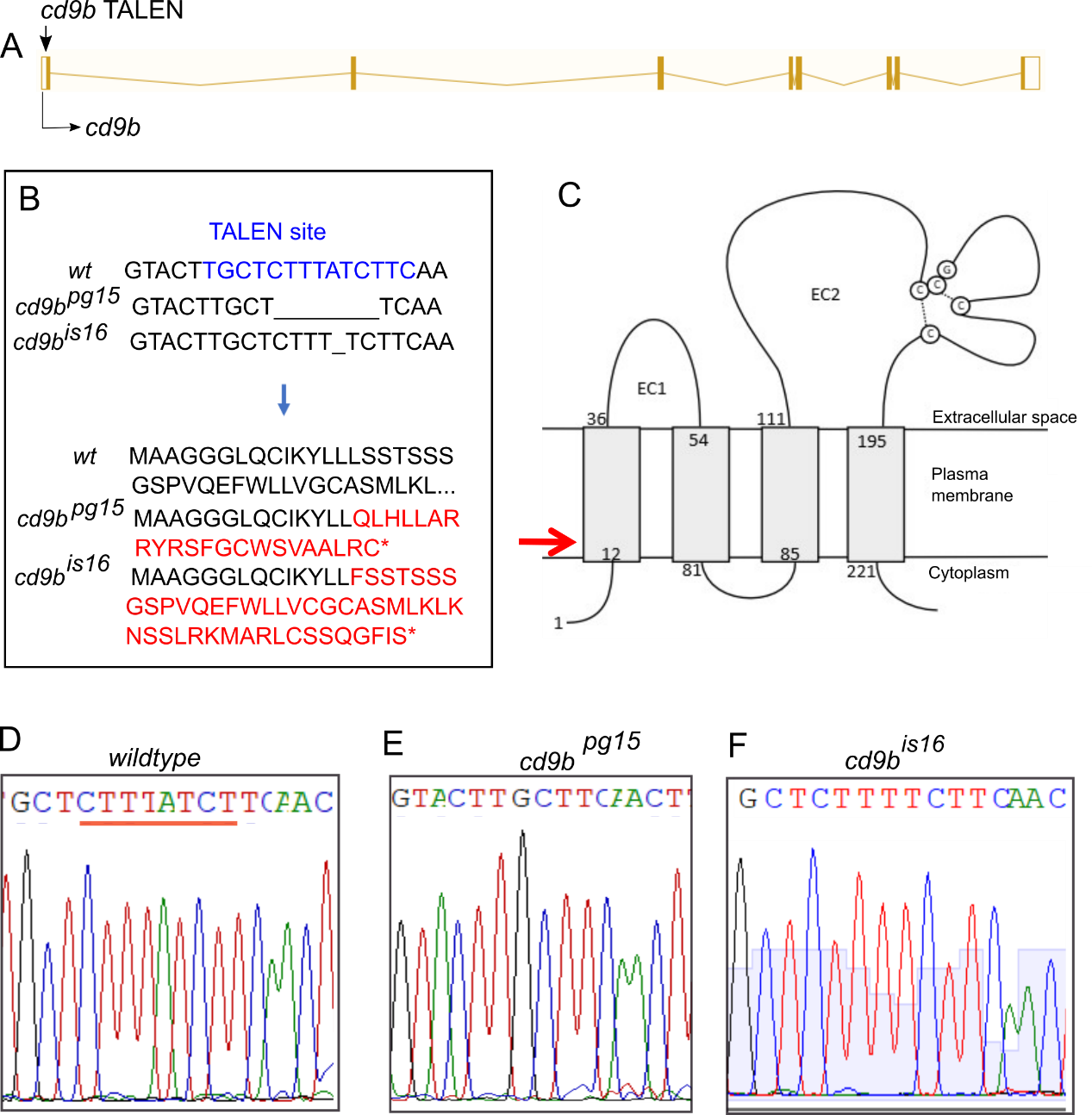
**Supplementary Figure 1. *cd9b* mutant generation.**

A: Nature of the *cd9b* mutant allele showing TALEN site location within the intron-exon structure of the gene. B: The TALEN target sequence in exon 1 is shown in blue; the 8bp deletion in the *cd9b^pg15^* allele, or the 1bp deletion in the *cd9b^is16^* allele is indicated under the WT sequence as dashes. The 8bp deletion leads to a frameshift changing codon 15 from TTT (Phe) to CAA (Glu), then 22 aberrant amino acids (red lettering) followed by a stop codon (*). The 1bp deletion leads to a frameshift changing codon 16 from ATC (Ile) to TCT (Ser), then 46 aberrant amino acids (red lettering) followed by a stop codon (*). C: Schematic of the Cd9b protein with location of mutation given by red arrow. The disulfide bonds between the conserved CCG motif and conserved cysteines are indicated by the dashed lines. EC1/2= Extracellular domain 1/2, aa= amino acid. D-F: Sequence chromatograms of genomic DNA from (d) WT and (e) *cd9b^pg15^* alleles and (f) *cd9b^is16^* alleles. Location of mutation is underlined in red.
